# Quercetin selectively reduces expanded repeat RNA levels in models of myotonic dystrophy

**DOI:** 10.1101/2023.02.02.526846

**Authors:** Subodh K. Mishra, Sawyer M. Hicks, Jesus A. Frias, Sweta Vangaveti, Masayuki Nakamori, John D. Cleary, Kaalak Reddy, J. Andrew Berglund

## Abstract

Myotonic dystrophy is a multisystemic neuromuscular disease caused by either a CTG repeat expansion in *DMPK* (DM1) or a CCTG repeat expansion in *CNBP* (DM2). Transcription of the expanded alleles produces toxic gain-of-function RNA that sequester the MBNL family of alternative splicing regulators into ribonuclear foci, leading to pathogenic mis-splicing. There are currently no approved treatments that target the root cause of disease which is the production of the toxic expansion RNA molecules. In this study, using our previously established HeLa DM1 repeat selective screening platform, we identified the natural product quercetin as a selective modulator of toxic RNA levels. Quercetin treatment selectively reduced toxic RNA levels and rescued MBNL dependent mis-splicing in DM1 and DM2 patient derived cell lines and in the *HSA*^LR^ transgenic DM1 mouse model where rescue of myotonia was also observed. Based on our data and its safety profile for use in humans, we have identified quercetin as a priority disease-targeting therapeutic lead for clinical evaluation for the treatment of DM1 and DM2.

**One Sentence Summary:** The natural product quercetin reduces toxic RNA in myotonic dystrophy.

## INTRODUCTION

Expansions of microsatellite sequences in coding and non-coding regions of the human genome cause over 50 neurological, neurodegenerative, and neuromuscular diseases including myotonic dystrophy (DM) (*1*). DM is the most common form of adult-onset muscular dystrophy and is a complex multisystemic disorder characterized by myotonia, myopathy, cardiac conduction abnormalities, gastrointestinal issues, sleep disturbance, and cognitive disruption (*2*). There are 2 genetically distinct forms of DM: type 1 (DM1) is caused by a CTG repeat expansion in the 3’ untranslated region (UTR) of the dystrophia myotonica protein kinase (*DMPK*) gene (*3-5*) while type 2 (DM2) is caused by a CCTG repeat expansion in intron 1 of the CCHC-type zinc finger nucleic acid binding protein (*CNBP*) gene (*6*). While the unaffected population typically have fewer than 50 repeats within either locus, affected individuals can have several hundred to thousands of CTG or CCTG repeats (*7*). Global prevalence studies typically estimate incidence of DM to be ∼1 in 8000 individuals, however a recent New York state newborn bloodspot genetic screening study reported the presence of DM repeat expansions to be approximately 1 in every 2100 births, suggesting that DM could be underdiagnosed (*8*).

A major pathogenic mechanism in DM involves transcription of the expanded alleles producing CUG/CCUG gain-of-function toxic expansion RNA that sequester the muscleblind-like (MBNL) family of RNA processing factors into ribonuclear foci. This sequestration results in mis-splicing of developmentally regulated cassette exons, leading to a host of disease symptoms (*7*). For example, mis-splicing of the muscle-specific chloride channel (*CLCN1)* pre-mRNA leads to myotonia which can be rescued in a mouse model by inducing the correct splice isoform (*9*). For this reason, mis-splicing serves as an important biomarker of disease (*10, 11*) and as a tool for identifying and evaluating therapeutics in DM.

There are currently no disease targeting therapies for DM and the few treatments available are aimed at the management of a limited number of symptoms. Thus, there is a critical need for treatments targeting the root cause of DM which is the production of the toxic expansion RNA. Pre-clinical studies have established proof-of-concept for the efficacy of modulating the toxic RNA in DM1/2 using a variety of approaches including small molecules, antisense oligonucleotides, RNA interference and CRISPR/Cas9 [Reviewed in (*12-14*)]. However, none of these approaches have yet to achieve clinical success due to toxicity, limited bioavailability and tissue distribution of the therapeutic agent, and insufficient target engagement among other limitations.

Previously, we developed a DM1 HeLa CTG repeat-selective screening platform to facilitate the discovery of small molecules that selectively reduce the toxic CUG RNA in myotonic dystrophy type 1 (*15*). Here, using this assay, we conducted an unbiased screen of a natural compound library leading to the identification of the flavonoid centaureidin, which selectively reduced toxic CUG RNA levels. Subsequent targeted screening of representative compounds from the large flavonoid family identified a member of the flavonol sub-class, quercetin, as the leading candidate with a favorable safety profile. Quercetin selectively reduced toxic RNA levels and rescued DM-associated mis-splicing in DM1 and DM2 patient derived cells in the absence of cellular toxicity. Treatment of the *HSA*^LR^ transgenic DM1 mouse model with the bioavailable form of quercetin, enzymatically modified isoquercitrin (EMIQ), selectively reduced toxic CUG RNA levels, rescued DM-associated mis-splicing and reduced myotonia. Our findings establish quercetin as a strong disease-targeting therapeutic lead with a well-established safety profile, that is primed for evaluation in clinical trials for DM.

## RESULTS

### Screening of a natural product library identifies flavonoids as a novel class of compounds that selectively reduce toxic CUG RNA levels

To identify small molecules that selectively reduce toxic CUG RNA levels and with favorable toxicity profiles, we selected the Natural Products Set V library from the National Cancer Institute. This library, which consists of 390 diverse compounds, was screened using our established HeLa DM1 CTG repeat-selective screening platform (*15*). Cells were treated in 96-well format at a concentration of 1 µM for 24 hours, permeabilized, and RNA was taken directly forward to multiplex RT-qPCR using probes targeting r(CUG)480 expansion RNA levels relative to r(CUG)0 control RNA as done previously (*15*) (Fig. 1A). Each screening plate contained triplicates of a DMSO control, media only control and as positive controls, colchicine (1 µM) and actinomycin D (ActD, 20 nM) treatments. As previously observed (*15*), colchicine and ActD treatments yielded a selective reduction in r(CUG)480 levels of ∼ 0.5, which served as our hit threshold (fig. S1). We ranked each compound using a cut-off of < 0.5, which revealed 6 primary hits from the library (Fig. 1, B and C) - three hits were from the ansamycin family, two hits were flavonoids, and one hit belonged to the alkaloid family of small molecules.

**Fig. 1.**
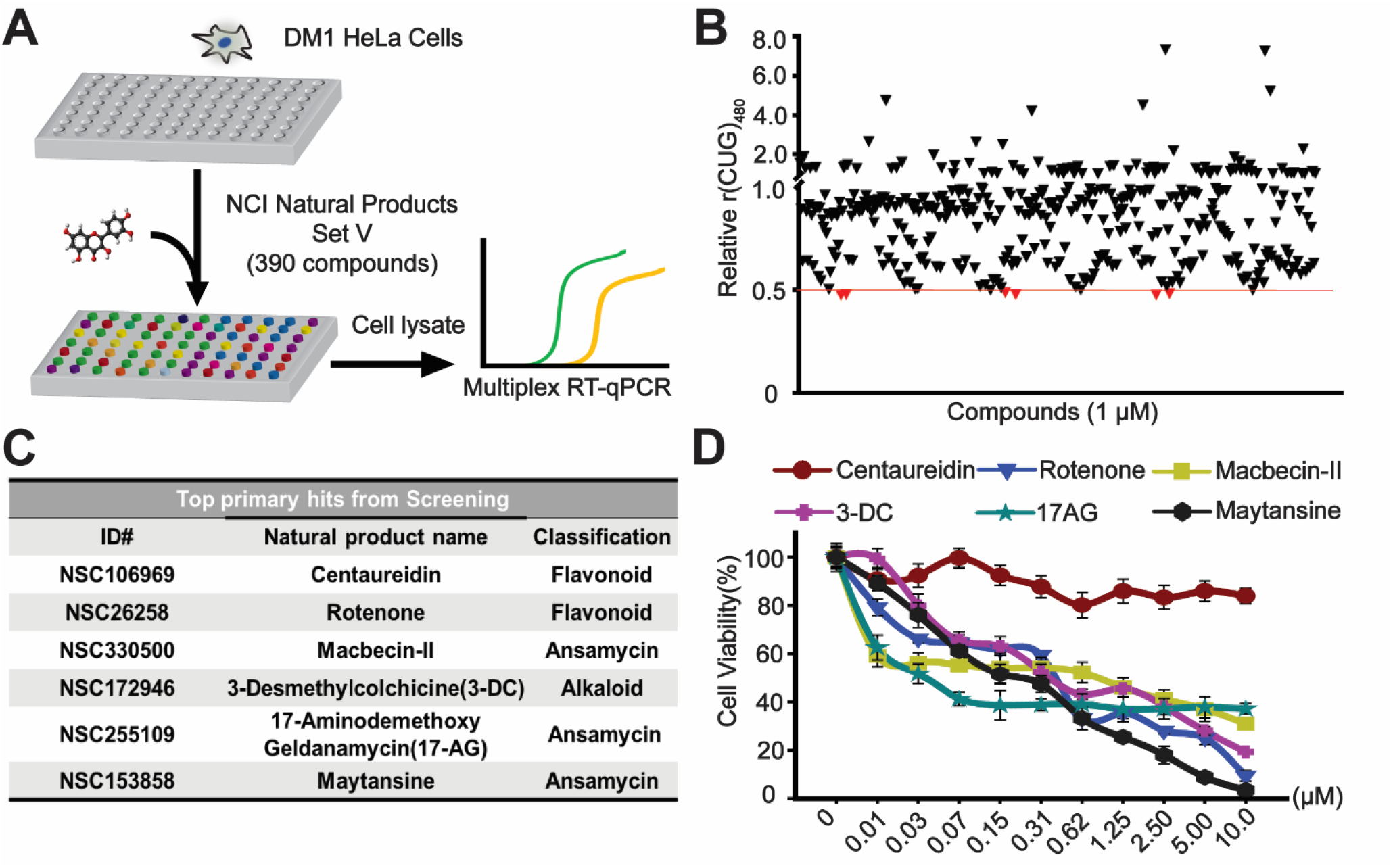
CTG repeat-selective HeLa DM1 cell screen of natural products. (**A**) Schematic of DM1 HeLa cellular screen of National Cancer Institute (NCI) Natural Product Set V (390 compounds). (**B**) Screening results of each drug treatment (1µM) on r(CUG)480 levels relative to r(CUG)0 levels normalized to DMSO treatment. The red line indicates the 0.5 cutoff to identify primary hits falling at or below the threshold (red dots are the 6 primary hits). (**C**) List of the 6 primary screening hits and natural product classification. (**D**) MTT cell viability assay conducted on HeLa DM1 cells to compare the relative toxicity of the 6 primary hits treated for 24 hours at the indicated concentration, normalized to DMSO treatment (mean ± SD, n= 3 biological replicates).

To prioritize hits with the lowest cellular toxicity for further testing, we next performed MTT assays which measure NAD(P)H-dependent oxidoreductase enzyme activity as an indicator of cell viability, proliferation, and cytotoxicity. Among the 6 lead hits, the flavonoid centaureidin was found to display the least amount of toxicity in the DM1 HeLa cell model across a broad dose range (0.01 - 10 µM, Fig. 1D). Flavonoids are a class of polyphenolic natural phytocompounds that are abundantly present in various flowers, fruits and vegetables and thus are normally consumed as part of our daily diet (*16*). Flavonoids can be broadly categorized into eight subclasses based on several factors such as degree of hydroxylation, degree of methylation, glycosylation pattern, degree of oxidation, degree of unsaturation, and the position of the carbon of the C ring to which the B ring is attached (Fig. 2A) (*17*). Given the large structural diversity within the flavonoid family, we selected several representative molecules from each subclass and tested their capacity to selectively down-regulate r(CUG)480 levels in the DM1 HeLa cell model (Fig. 2B). Ranking compounds based on the degree of the selective reduction in r(CUG)480 levels revealed that the flavonol subclass which includes the primary hit centaureidin, contained the most active hits (Fig. 2B, red bars). Among the flavonols, centaureidin and quercetin showed the greatest selective reduction of r(CUG)480 levels at the screening dose of 1µM (Fig. 2B). However, quercetin exhibited lower cellular toxicity compared to centaureidin in the DM1 HeLa cell model positioning it as our top lead (Fig. 2C).

**Fig. 2.**
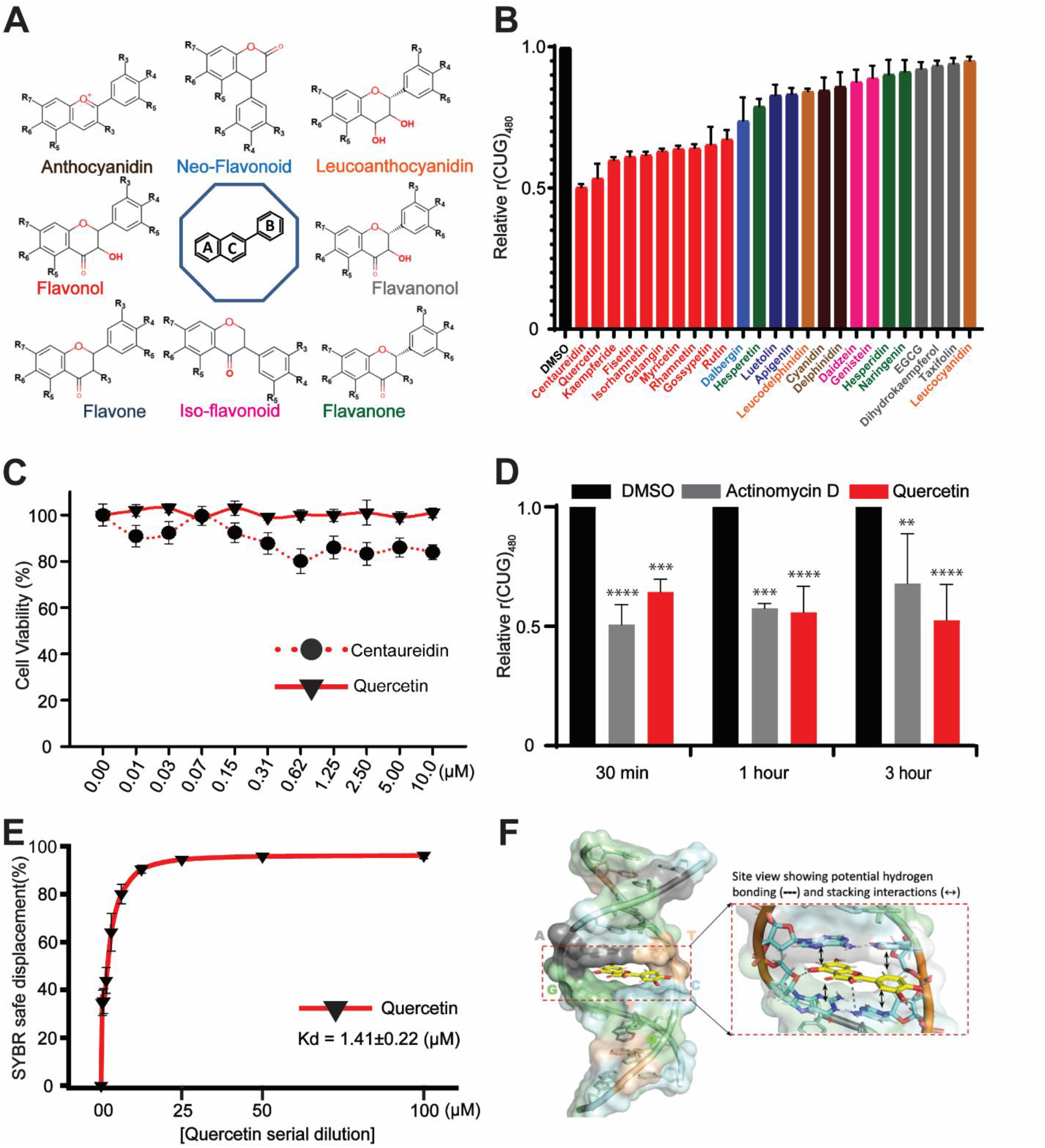
Targeted flavonoid screen identifies quercetin as a selective modulator of toxic CUG RNA levels. (**A**) Structures of the various sub-classes of flavonoids. R denotes positions of the various functional groups within each sub-class. (**B**) Targeted screen of selected flavonoids from each sub-class in HeLa DM1 cells treated for 24 hours at 1µM. r(CUG)480 levels relative to r(CUG)0 normalized to DMSO treatment (mean ± SD, n= 3 biological replicates). (**C**) MTT viability assay conducted on HeLa DM1 cells treated with centaureidin or quercetin for 24 hours, normalized to DMSO treatment (mean ± SD, n= 3 biological replicates). (**D**) 5-ethynyl uridine (EU) nascent RNA labeling assay. HeLa DM1 cells were treated with quercetin or actinomycin D for 24 hours then transcription was blocked with DRB (5,6-dichlorobenzimidazole 1-β-d-ribofuranoside) for 3.5 hours, followed by washing out DRB and labeling with EU in the presence of quercetin (1µM) or Actinomycin D (ActD) (20 nM) for the indicated times. Multiplex RT-qPCR was performed on RNA labeled with EU to determine nascent RNA levels of r(CUG)480 relative to r(CUG)0 normalized to DMSO treatment] (mean ± SD, n = 3 biological replicates). (**E**) Fluorescent intercalator displacement (FID) assay on a (CTG)2(CAG)2 palindromic duplex substrate bound by SYBR Safe fluorescent probe in the presence of increasing quercetin (0.39 µM - 100 µM) compared to DNA template without quercetin (0 µM) (mean ± SD, n = 3 technical replicates). (**F**) Computational modeling of a 10-mer (CTG)·(CAG) repeat DNA with quercetin intercalating at a CT·AG site. The magnified site view highlights hydrogen bonding (dotted lines) and stacking interactions (arrows) within the binding pocket.

The reduction in toxic CUG RNA levels in our screening assay could be the result of reduced transcription or of increased RNA turn-over. To distinguish between these two effects, we carried out 5-ethynyl uridine (EU) pulse labeling and nascent RNA isolation followed by multiplex RT-qPCR for relative r(CUG)480 and r(CUG)0 levels in the HeLa DM1 cells as done previously (*15*) (Fig. 2D). ActD, a transcriptional inhibitor, which was previously shown to selectively reduce transcription of CTG expansions at low doses (*18*), served as a positive control. Similar to ActD treatment, varying EU pulse times between 30 min and 3 hours did not significantly alter the selective reduction in r(CUG)480 levels following quercetin treatment (Fig. 2D). This finding is consistent with selective transcription inhibition of the CTG expansion by quercetin. Previous studies have reported the DNA intercalation potential of quercetin and related flavonoids (*19, 20*). Accordingly, fluorescent intercalator displacement (FID) assay confirmed that quercetin was able to competitively displace bound SYBR green probe from a (CTG)2(CAG)2 duplex substrate *in vitro* with a calculated K_D_ of 1.41 µM (Fig. 2E). There are three unique intercalation sites within the (CTG)·(CAG) DNA repeat based on the flanking nucleotide composition. *In silico* molecular modeling and docking predicts intercalation at the (CT)·(AG) site to be more thermodynamically favorable than the others (Fig. 2F). At this site, The polycyclic benzopyran ring of quercetin stacks with the purines and the phenyl ring stacks with the pyrimidines in a canonical purine:pyrimidine base-pair like configuration (Fig 2F). In addition, the hydroxyl groups of quercetin also form several hydrogen bonds with the backbone on either side of the intercalation site further anchoring it in place. While similar stacking interactions are also observed in the predicted complexes of the other two intercalation sites, the primary stacking interactions in those cases involve only the benzopyran ring and one of the flanking purines. Overall, the docking predictions indicate that quercetin has the right size and can assume a conformation that is suitable for intercalation with CTG repeats. While our *in silico* model and FID assay supports favorable binding of quercetin to the (CTG)·(CAG) DNA helix through intercalation, quercetin could also potentially bind multiple sites within the expanded repeat and/or through other binding modes. In summary, our data is consistent with selective transcriptional inhibition of the CTG expansion through quercetin directly binding the (CTG)·(CAG) repeats.

In summary, our unbiased natural product screen and subsequent targeted flavonoid screen identified quercetin as the top hit based on the selective reduction of r(CUG)480 levels and minimal cellular toxicity in the HeLa DM1 cell model. The United States Food and Drug Administration (FDA) designates quercetin as generally recognized as safe (GRAS) based on the low toxicity and general tolerance in humans (*21*). Taken together, this data supported the prioritization of quercetin as the lead therapeutic candidate for additional evaluation in multiple DM models.

### Quercetin rescues mis-splicing in DM patient-derived fibroblasts

We validated the effect of quercetin treatment on mis-splicing in female and male DM1 patient derived fibroblast cell lines (DM1-A and DM1-B respectively). Cells were treated for 24 hours at a dose range between 0.5 µM to 128 µM, followed by RNA extraction and RT-PCR analysis for two MBNL dependent mis-splicing events, *INSR exon 11* and *FLNB exon 31*. Quercetin treatment resulted in a significant dose dependent rescue of mis-splicing between 16 – 128 µM with ∼100% splicing rescue of both events at 64 µM in both DM1-A (Fig. 3A) and DM1-B (fig. S2A) cell lines. Treatment of unaffected control fibroblasts did not result in significant alterations to the splicing outcome of the same events (fig. S2B). Based on the shared mechanism of toxic RNA pathogenesis in DM1 and DM2, we also tested the effect of quercetin treatment in characterized DM2 patient derived fibroblasts (*22*) using the same mis-splicing events (fig. S3A). A comparable rescue of mis-splicing was observed in the DM2 cell line as was observed in DM1 cell lines (fig. S3A). Because RNA toxicity stems from production of CUG and CCUG expansion RNA from the *DMPK* and *CNBP* genes respectively, we evaluated the effect of quercetin treatment on expression of *DMPK* in DM1 treated cells (Fig. 3B and fig. S2C) and *CNBP* in DM2 treated cells (fig. S3B). There was a significant dose-dependent reduction in both *DMPK* and *CNBP* genes in DM1 (Fig. 3B and fig. S2C) and DM2 fibroblasts (fig. S3B) respectively but not in unaffected controls (Fig. 3B and fig. S3B), consistent with an effect mediated through the toxic RNA. It should be noted that in this assay we are not distinguishing the expanded from the unexpanded alleles. Consistent with findings from the HeLa DM1 cell model, treatment of fibroblast cell lines with quercetin across a broad dose range (0.5 to 128 µM) revealed negligible toxicity (MTT assay) in DM and unaffected control cell lines (Fig. 3C, fig. S2D and S3C).

**Fig. 3.**
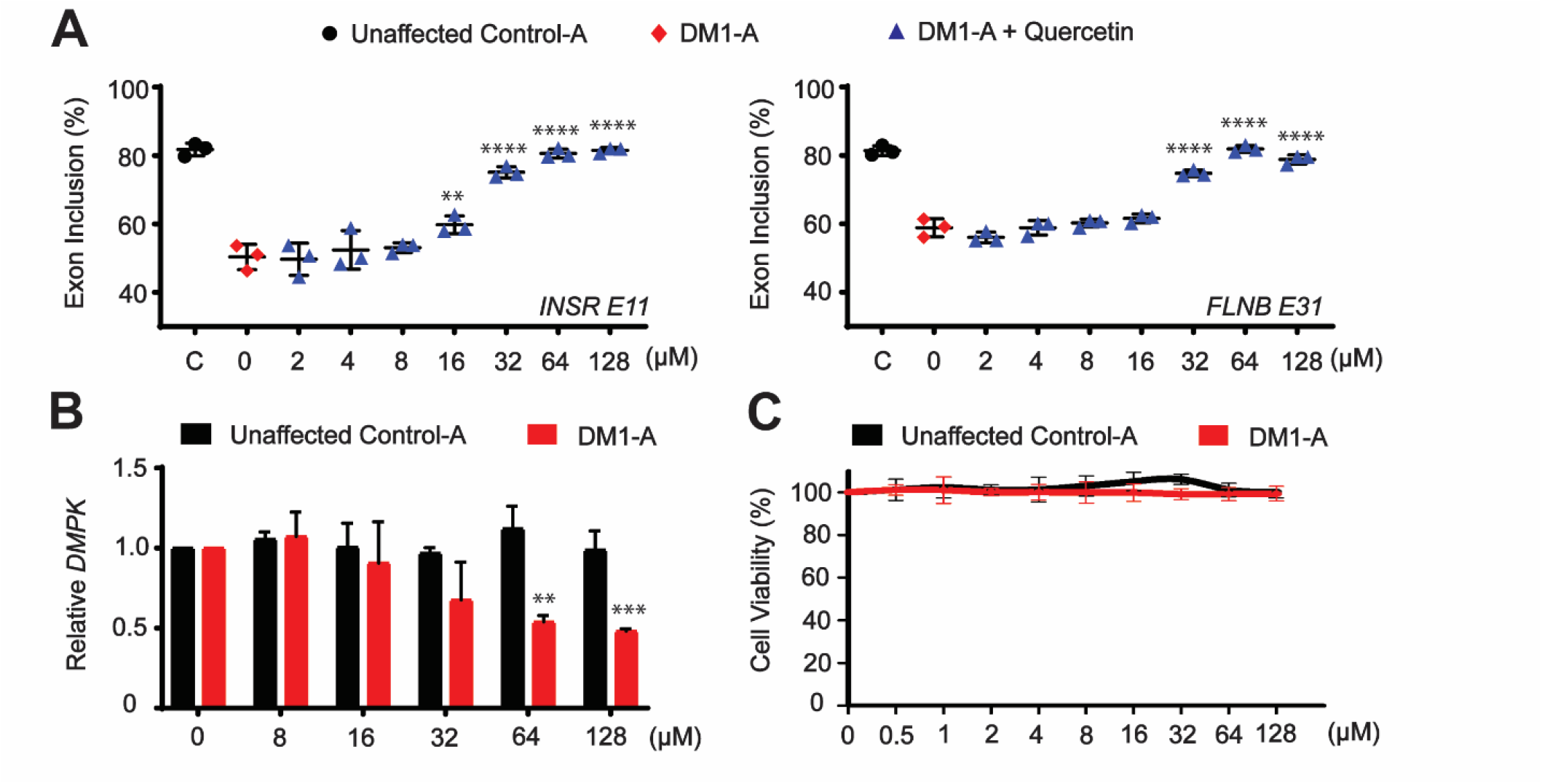
Validation of quercetin activity in DM1 patient derived fibroblasts. (**A**) RT-PCR cassette exon isoform analysis of *INSR exon 11* (left) and *FLNB exon 31* (right) alternative splicing events following 24-hour treatment with quercetin (Mean ± SD *n* = 3 biological replicates). (**B**) RT-qPCR analysis of *DMPK* levels (relative to *GAPDH*) following quercetin treatment normalized to DMSO control in DM1 and unaffected control fibroblasts (Mean ± SD *n* = 3 biological replicates). (**C**) Cell viability assay performed on unaffected control and DM1 fibroblasts following 24-hour treatment with quercetin at the indicated concentrations. Normalized to DMSO control (0 µM) (Mean ± SD, n = 3 biological replicates). One-way ANOVA with Dunnett’s multiple comparisons test comparing treated fibroblasts to DMSO control \*\**P* <0.01, \*\*\**P* < 0.001 \*\*\*\**P* < 0.0001.

### Quercetin reduces toxic RNA levels and rescues DM-associated mis-splicing independent of senolytic activity

Quercetin and other flavonoids including fisetin have been described as senolytics, compounds that selectively induce cell death in senescent cells (*23*). Recent studies demonstrated that accelerated senescence, which is observed in myotonic dystrophy models, could be modulated using senolytics (*24, 25*). In these studies, senolytic treatment reduced key molecular markers of senescence including p16 in cellular and animal models of DM1, supporting senolytics as potential therapeutics for DM (*24, 25*). However, the effects of quercetin and/or other senolytics on key molecular hallmarks of RNA toxicity including mis-splicing were not determined (*24, 25*). To test if senolytic activity contributes to toxic RNA-mediated mis-splicing, we compared quercetin to the non-flavonoid senolytics, dasatanib and A1155463 in DM1 patient fibroblasts (fig. S4). Consistent with previous findings using senolytics, all compounds reduced the p16 marker of senescence that was significantly elevated in DM1 cells (fig. S4A). However, in contrast to the reduction of *DMPK* levels and rescue of mis-splicing in DM1 patient fibroblasts by quercetin, treatment with dasatinib or A1155463 did not affect *DMPK* levels or rescue mis-splicing (fig. S4B and C). Thus, while senolytics may represent an important therapeutic approach for DM through modulating accelerated senescence, quercetin rescues toxic RNA mediated mis-splicing independent of senolytic activity.

### Validation of quercetin activity in DM1 patient-derived myotubes

Muscles are one of the primary affected tissues in DM and thus we sought to test the effects of quercetin using patient derived myoblasts. Myoblasts from a DM1 patient containing an expansion of ∼1,900–3,000 CTG repeats in the 3′ UTR of *DMPK (26)* were differentiated into post mitotic myotubes, treated with quercetin and then evaluated for MBNL dependent missplicing. DM1 myotubes treated with quercetin for 72 hours revealed a dose-dependent rescue of *MBNL1* exon 5 and *SYNE1* exon 137 with a maximal rescue at 32 µM (Fig. 4A). Interestingly, full rescue was observed for *SYNE1* exon 137 and only a partial rescue of *MBNL1* exon 5 (∼ 40 %) at 32 µM. Notably, there was reversion in the splicing rescue at and beyond 64 µM for both *MBNL1* and *SYNE1* events, concurrent with a decrease in cellular viability at those higher doses (Fig. 4B). Treatment of unaffected control myotubes performed in parallel did not reveal changes to the splicing profile of the same events nor did treatment of DM1 myotubes reveal changes to previously identified (*22*) non-DM1 associated alternative splicing events (fig. S5, S6). Consistent with an effect mediated through the toxic CUG RNA, treatment of DM1 myotubes with quercetin significantly reduced overall *DMPK* expression whereas *DMPK* levels were not affected in control myotubes (Fig. 4C). Taken together with the fibroblast results, quercetin treatment reduces *DMPK* levels and rescues DM-associated mis-splicing, a key molecular hallmark of toxic RNA, in patient derived cell lines with marginal effects on cellular toxicity within the dose range showing optimal rescue.

**Fig. 4.**
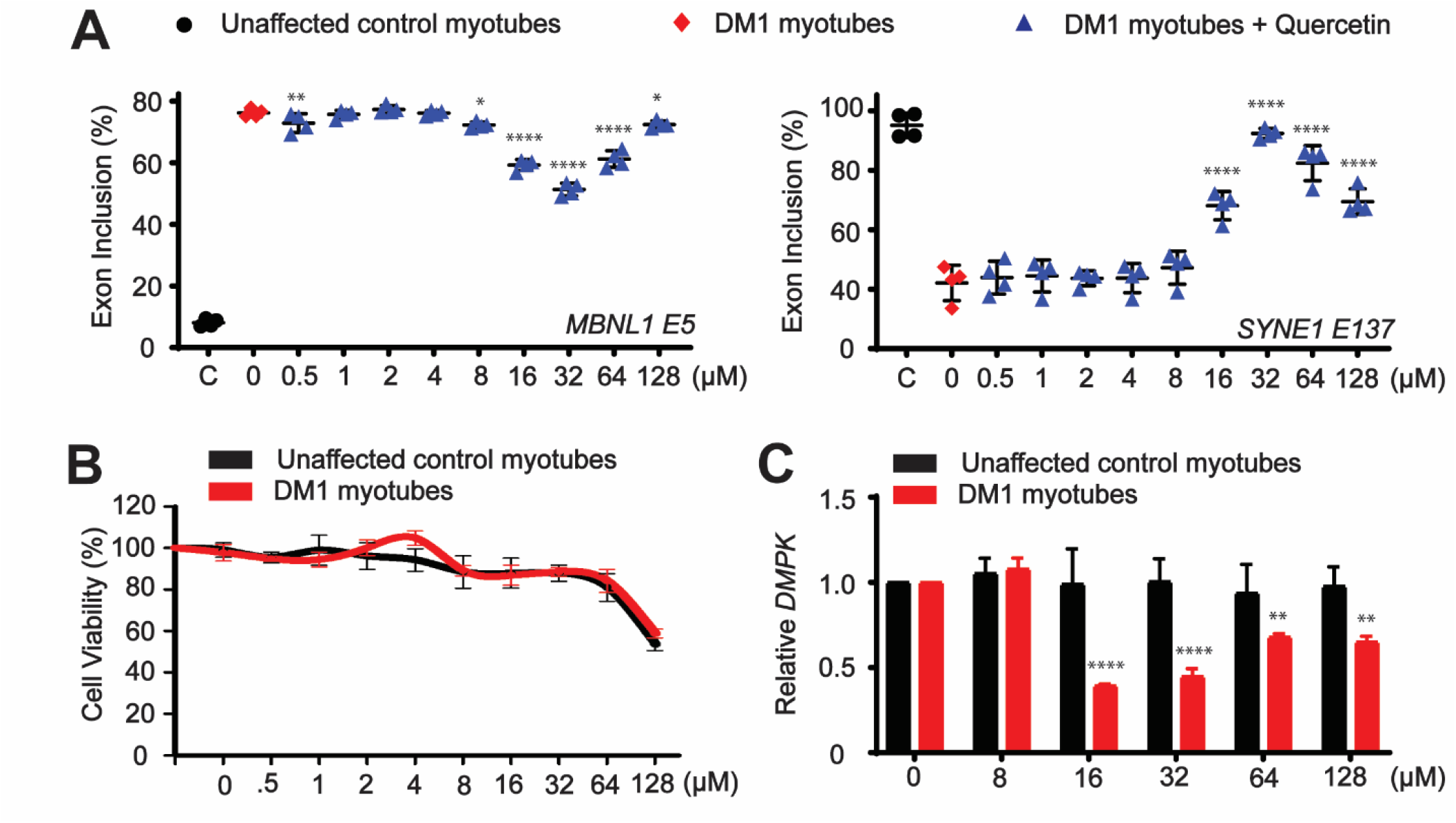
Validation of quercetin activity in DM1 patient derived myotubes. (**A**) RT-PCR cassette exon isoform analysis of the *MBNL1 exon 5* (left) and *SYNE1 exon 137* (right) alternative splicing events following 72-hour treatment with quercetin (Mean ± SD *n* = 4 biological replicates). (**B**) Cell viability assay performed on unaffected control and DM1 myotubes following 72-hour treatment with quercetin at the indicated concentrations. Normalized to DMSO control (µM) (Mean ± SD, n = 3 biological replicates). (**C**) RT-qPCR analysis of *DMPK* levels (relative to *GAPDH*) following quercetin treatment in DM1 and unaffected control myotubes normalized to DMSO control (0µM) (Mean ± SD *n* = 4 biological replicates). One-way ANOVA with Dunnett’s multiple comparisons test comparing treated myotubes to DMSO control (0 µM) \**P* < 0.05, \*\**P* < 0.01, \*\*\**P* < 0.001, \*\*\*\**P* < 0.0001.

### Treatment of DM1 *HSA*^LR^ mice with a quercetin derivative reduces toxic CUG RNA levels, rescues DM-associated mis-splicing and improves myotonia

To determine the *in vivo* efficacy of quercetin, we investigated quercetin activity in the DM1 *HSA*^LR^ transgenic mouse model (*27*). *HSA*^LR^ mice express a human skeletal actin (*HSA)* transgene containing ∼220 CTG repeats within an untranslated region of the final exon, in the absence of *DMPK* sequence context (*27*). Given that quercetin has a well-established solubility issue *in vivo*, we evaluated enzymatically modified isoquercitrin (EMIQ), a glucoside derivative of quercetin with improved solubility and bioavailability (fig. S7) (*28-31*). EMIQ, like unmodified quercetin, is also recognized by the FDA as safe (GRAS) and is widely sold as a nutritional supplement. EMIQ was administered daily to *HSA*^LR^ mice through their drinking water at low (1.5 g/L) or high (15 g/L) doses, for 6 and 12 weeks. Following treatment, total RNA was extracted from quadriceps muscle and analyzed using RT-qPCR to assess changes in gene expression and splicing (Fig. 5). While low dose EMIQ treatment was insufficient to reduce *HSA* transgene expression (Fig. 5A), both 6 and 12-week high dose treatments resulted in a significant reduction in *HSA* transgene mRNA containing the CUG expansion (Fig. 5A). There was no significant change to expression of endogenous *Dmpk* which does not contain a CUG repeat tract (Fig. 5A). Interestingly, a shorter 3-week high dose EMIQ treatment was insufficient to reduce *HSA* transgene mRNA suggesting that the effect may have latency (fig. S8). Selectively reducing *HSA* CUG transgene expression is expected to rescue MBNL dependent mis-splicing and thus we carried out RT-PCR analysis of established DM-associated alternative splicing events. Treatment of *HSA*^LR^ mice with a high dose of EMIQ for either 6 or 12 weeks resulted in a significant and comparable rescue of *Clcn1* exon 7 and *Atp2a1* exon 22 mis-splicing (Fig. 5B). As observed with *HSA* transgene expression, low dose EMIQ treatments did not show splicing recue following either 6-or 12-week treatments, nor did high dose treatment for 3 weeks (Fig. 5B, fig. S8).

**Fig. 5.**
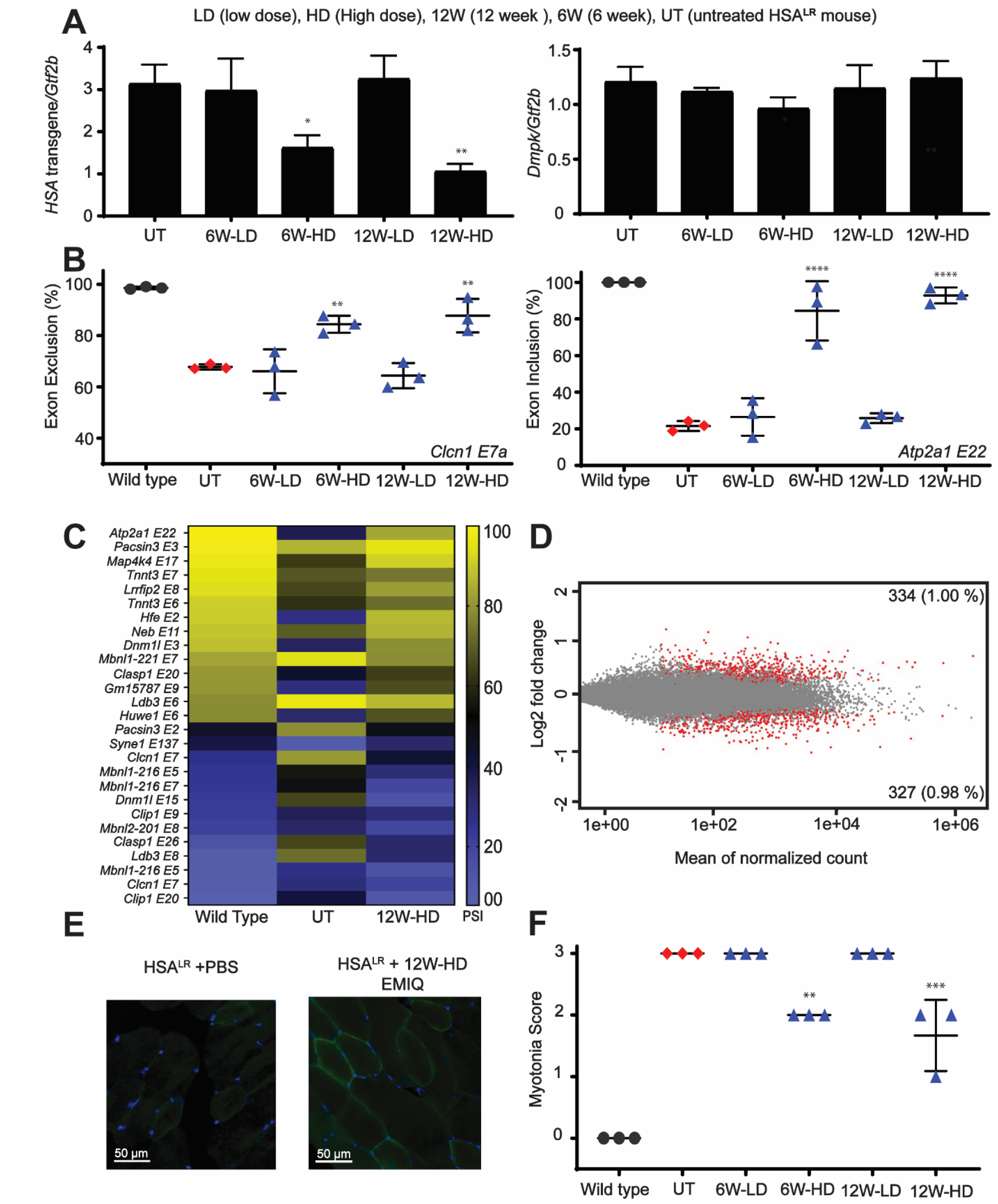
Rescue of splicing and myotonia in the DM1 *HSA*^LR^ mouse model. (**A**) RT-qPCR analysis of *HSA* transgene (left panel) or endogenous *Dmpk* (right panel) expression (normalized to *Gtf2b*) following EMIQ administration in the drinking water for 6 weeks (6W) or 12 weeks (12W) at a low dose of 1.5 g/L (LD) or high dose of 15 g/L (HD). (Mean ± SD, n = 3 biological replicates), \**P* < 0.05, \*\**P* < 0.01. (**B**) RT RT-PCR cassette exon isoform analysis of *Clcn1 exon 7a* (left) and *Atp2a1 exon 22* (right) alternative splicing events following EMIQ administration in the drinking water at the indicated concentrations (Mean, n = 3, *P < 0.05, **P < .01). (**C**) Heat map displaying the rescue of alternative cassette exon inclusion by mean normalized PSI identified from the top 100 mis-splicing events in 12-week high dose EMIQ treated *HSA*^LR^ mice compared to *HSA*^LR^ mice without treatment and wild-type mice (*P* < 0.0005, FDR < 0.05). (**D**) MA plot displaying differential gene expression comparing untreated *HSA*^LR^ mice vs. *HSA*^LR^ mice treated with high-dose EMIQ for 12-weeks. Log2 fold gene expression changes are plotted by mean normalized counts. Red dots represent changes with adjusted *P* < 0.1. Grey dots represent non-significant events. (**E**) Immunofluorescence detecting Clcn1 protein in 12-week high dose EMIQ treated and untreated *HSA*^LR^ mouse quadriceps muscle sections. Nuclei are stained using DAPI and represented in blue. (**F**) Comparison of myotonia grade using electromyography needle insertions between wild type, untreated *HSA*^LR^ mice and *HSA*^LR^ mice treated with 12-week high dose of EMIQ (Mean, n = 3, *P < 0.05, **P < .01).

To broadly evaluate transcriptome alterations resulting from EMIQ treatment, we carried out RNAseq analysis from quadriceps muscle of *HSA*^LR^ mice treated with high dose EMIQ for 12 weeks. This treatment displayed the greatest reduction of *HSA* transgene mRNA and rescue of candidate mis-splicing (Fig. 5C, D). Using the RNAseq data we evaluated a panel of cassette exon alternative splicing events that showed the greatest percent spliced in (PSI) difference between wild-type and *HSA*^LR^ mice. This panel included well characterized mis-spliced pre-mRNAs contributing to muscle phenotypes in DM1 such as *Atp2a1, Clcn1, Clasp1, Syne1*, and *Tnnt3 (10, 11)*. We observed broad rescue across this panel with multiple targets approaching complete rescue (Fig. 5C and data table S1). Gene expression analysis revealed modest changes to the transcriptome following 12-week EMIQ high dose treatment in *HSA*^LR^ mice with ∼2% of the transcriptome significantly altered, and only two genes showing greater than 2-fold expression changes (Fig. 5D and data table S2). It should be noted that these alterations are likely an overestimate of off-targets because toxic CUG RNA expression is known to deregulate gene expression (*32*) and thus some of these significant changes may reflect rescue in gene expression following EMIQ treatment. These data indicate that EMIQ selectively reduces expanded CUG RNA levels and reverses DM-associated mis-splicing without broadly disrupting the transcriptome.

The RNAseq analysis supported broad reversal of multiple pathogenic mis-splicing events, including *Clcn1* exon7a which results in reduced Clcn1 protein leading to myotonia in DM (*9, 33*). Immunofluorescence staining for Clcn1 protein in quadriceps muscle sections from *HSA*^LR^ mice treated with high dose EMIQ for 12 weeks confirmed increased Clcn1 expression compared to untreated *HSA*^LR^ mice (Fig. 5E). To understand the functional outcomes of EMIQ treatment, we examined myotonia using electromyography (EMG) analysis of treated animals. A graded myotonia score of 3 indicated frequent repetitive discharges in quadriceps muscle for nearly all electrode insertions in untreated *HSA*^LR^ mice. Treatment with high dose EMIQ for 6 or 12 weeks resulted in improved myotonia scores of 2 and 1.5 respectively (Fig. 5F). Collectively, these results demonstrate that *in vivo* EMIQ treatment administered through drinking water selectively reduces toxic CUG RNA levels, resulting in reversal of mis-splicing and functional rescue of disease phenotypes associated with DM1 in a mouse model.

## DISCUSSION

Using our previously established DM1 HeLa repeat selective screening platform (*15*), we identified flavonoids as a novel class of natural compounds that selectively reduce toxic CUG RNA levels. Subsequent targeted screening identified the flavonol sub-class to be the most active with quercetin emerging as the top hit. Quercetin was validated in DM1 and DM2 patient derived fibroblasts and in DM1 myotubes, revealing robust reversal of MBNL dependent mis-splicing without any appreciable cellular toxicity. Treatment of the *HSA*^LR^ DM1 mouse model with the orally bioavailable quercetin derivative EMIQ, validated its therapeutic potential *in vivo*, demonstrating robust reduction of toxic CUG RNA levels, reversal of DM associated mis-splicing and reduced myotonia. Our results of a disease targeting effect without inducing cytotoxicity, coupled with GRAS status, positions quercetin for future clinical evaluation for the treatment of DM.

Flavonoids have been previously tested in a PC12 neuronal cell model containing a (CTG)250 expansion and were found to reduce cytotoxicity (*34*). The mechanism of action however was not determined, and the effects of flavonoids on toxic CUG RNA pathogenesis were not tested. More recently, quercetin was also evaluated as a senolytic for treatment of DM1 (*24, 25*). In these studies, DM1 cells were reported to express the senescence associated secretory phenotype (SASP) and could be targeted by quercetin alone (*25*) or in combination with dasatinib (*24*). Both treatments lead to the induction of cell death in senescent DM1 cells. Quercetin treatment was also determined to prolong lifespan in a DM1 *Drosophila* model (*25*). While a senolytic effect was reported to be of therapeutic benefit in DM by reducing the senescent cell population and improving myogenic potential, the effects of quercetin and other senolytics on the molecular hallmarks of toxic CUG RNA including DM associated MBNL dependent mis-splicing were not reported. Our results establish a disease targeting effect of quercetin through reduction of toxic CUG RNA levels and rescue of downstream consequences including MBNL dependent mis-splicing and myotonia. Parallel comparison of non-flavonoid senolytics dasatinib and A1155463 did not reveal effects on DM-associated mis-splicing despite reduction of the senescence marker p16. Therefore we conclude that quercetin treatment modulates toxic CUG RNA levels directly and independently from its role as a senolytic. However, because quercetin treatment also led to a reduction in the elevated senescence-associated p16 marker in DM1 cells, consistent with previous studies demonstrating senolytic activity of quercetin in DM1 (*24, 25*), we propose that quercetin may act on at least two relevant therapeutic pathways in DM.

Selective reduction of expanded CUG/CCUG RNA by quercetin does not appear to be affected by the length of the CTG/CCTG repeats. The CTG and CCTG repeats tested in the cell lines and *HSA*^LR^ mouse model ranged from ∼220 CTG repeats in the *HSA*^LR^ mouse model (*27*) to ∼2,900 CTG repeats in DM1 myotubes (*26*) and ∼3000 CCTG repeats in DM2 fibroblasts (*22*). Additionally, the selective reduction of expansion RNA did not appear to be affected by genomic location of the repeat, which is present at different genomic loci in the HeLa DM1 cells (randomly integrated), DM1 (*DMPK*) and DM2 (*CNBP*) patient derived cell lines and in the *HSA*^LR^ mouse model (human skeletal actin transgene). The degree of splicing rescue by quercetin showed some variation across these systems. In fibroblasts, full splicing rescue was evident at 64 µM quercetin treatment, commensurate with reduced *DMPK* expression within the active dose range but no significant change in cellular viability (Fig 3). In DM1 myotubes however, quercetin treatment fully rescued mis-splicing of *SYNE1* exon 137 and only partially rescued *MBNL1* exon 5 with optimal rescue at 32 µM. Reduction of *DMPK* transcript was greatest at 16 and 32 µM and cell viability, which trended down over the quercetin concentration range, was reduced to 60% at 128 µM. The changes in cell viability at the highest concentration of quercetin treatment may explain the concomitant reduction in splicing rescue observed for the events tested.

The relatively rapid rescue of mis-splicing in cells by quercetin contrasts the need for several weeks of EMIQ treatment in the *HSA*^LR^ mice before splicing rescue is observed. The latency in the latter system may be due to processing of EMIQ to quercetin, the buildup of quercetin metabolites in tissues, or another aspect of delivery or uptake. While subtle variations were observed between the various systems, quercetin generally demonstrated a broad ability to rescue mis-splicing with minimal toxicity and transcriptomic alterations, making it an ideal candidate for future clinical testing.

DM involves multiple detrimental consequences downstream of toxic RNA production outside of MBNL sequestration, such as hyperphosphorylation of CELF1 (CUGBP Elav-Like Family Member 1) protein leading to additional RNA processing defects (*35*) and Repeat-Associated Non-ATG (RAN) translation leading to production of toxic peptide repeats (*36, 37*). Targeting multiple toxic pathways using combination therapies is a potential strategy to enable additive and synergistic effects. Proof-of-concept for this combination approach has already been demonstrated, where a combination of furamidine and erythromycin resulted in greater rescue of mis-splicing at lower doses than individual treatments in DM1 cell and mouse models, supporting this approach (*38*). EMIQ/quercetin, with its safety profile, ease of use and mechanism of action, is a strong candidate for this approach. For example, the FDA-approved drug, metformin was recently shown to inhibit RAN translation by modulating the double-stranded RNA-dependent protein kinase (PKR) pathway which is activated by expanded structured RNAs (*39*). Combining quercetin, which reduces toxic RNA levels in DM, with metformin to reduce DM RAN proteins could have a synergistic effect on DM pathogenesis. Alternatively, combining quercetin with the FDA-approved antibiotic erythromycin, a compound that disrupts MBNL1/2 sequestration, could lead to a synergistic rescue of RNA toxicity (*38*).

DM is part of a larger family of over 50 microsatellite expansion disorders (*7*), which includes *C9orf72* amyotrophic lateral sclerosis-frontotemporal dementia (C9 ALS/FTD), Huntington disease (HD), and numerous spinocerebellar ataxias (SCAs). Our results, demonstrating an effect of quercetin in both DM1 (CTG) and DM2 (CCTG) patient fibroblasts, supports the potential for quercetin to modulate expression of other repeat expansions. Future studies are necessary to investigate the effects of quercetin on other repeat expansion motifs including the GGGGCC repeats of C9 ALS/FTD and the CAG repeats of HD and various SCAs among others. In summary, our study has found that quercetin, a safe natural product, selectively reduces the production of toxic expansion RNA, the root cause of DM, and thus strongly warrants clinical investigation of quercetin for the treatment of DM.

## MATERIALS AND METHODS

### NCI Natural Product Set V screen

The Natural Product Set V 390 compound library was obtained from the developmental therapeutic program (DTP) of the National Cancer Institute (NCI). To screen the library compounds, our previously published HeLa DM1 CTG repeat-selective screening platform (*15*) was employed. Briefly, approximately 1×10^4^ cells were seeded in 96 well cell culture plates and cultured overnight in Dulbecco’s modified Eagle medium (DMEM) supplemented with 10% fetal bovine serum (FBS) and 1% penicillin and streptomycin under standard conditions of 37 °C and 5% CO_2_. The next day, the culture medium was replaced with fresh medium containing library compound or positive (1 µM colchicine or 20 nM ActD) or negative (DMSO) controls. Following 24 hours of treatment, each well of the treatment plates was washed with PBS and the plates were stored at - 80°C overnight. The next day, plates were thawed, and each well was treated with 20 µL of ‘lysis’ buffer (0.25% Igepal buffer, 10 mM HCl pH = 7.5, and 150 mM NaCl) on an orbital shaker for 5 min at room temperature. Then, 2 µL of cell ‘lysate’ was directly used for cDNA generation using HT_RT primer (table S1) and SuperScript IV reverse transcriptase (Thermo Fisher). Following cDNA synthesis, multiplex qPCR was performed on a Bio-Rad C1000 Touch Thermal Cycler using Hot Start Taq 2× Master Mix (NEB) with HT_Forward and HT_Reverse primers (IDT) (table S1) and custom HT_Probe1 and HT_Probe2 fluorescent probes (IDT) (table S1). Data were analyzed using the comparative (ΔΔCT) method. The levels of r(CUG)480 relative to r(CUG)0 were normalized to DMSO control treatments.

### Cell culture toxicity analysis

Cytotoxicity of Quercetin and other natural compounds was evaluated using MTT (3-(4, 5- dimethylthiazol-2-yl)-2,5 diphenyltetrazolium bromide dye) assay (Sigma Aldrich). Following the drug treatment in triplicate at indicated concentrations, cells were washed with PBS and replaced with a 1:1 ratio of fresh growth medium and 5mg/mL of MTT (Sigma Aldrich) stock solution prepared in 1x PBS. Next, MTT-containing cells were grown for additional 4 hours at 37°C and 5% CO_2_ to allow the intracellular reduction of the soluble yellow MTT to insoluble formazan crystals, followed by the addition of 100 µL of DMSO at Room temperature and shaking at orbital shaker for 5 minutes. DMSO dissolved the formazan crystal and absorbance was measured at 590 nm on a microplate reader (Synergy™ H1 multi-mode microplate reader). The percentage of living cells was calculated by using equation 2. Where ʎ590 is absorbance measured at 590 nm.

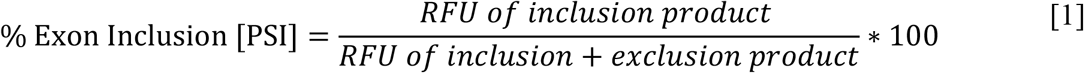

### Ethynyl uridine pulse labeling and nascent RNA isolation

Protocol was carried out as described previously (*15*). Briefly, approximately 1×10^5^ DM1 HeLa cells were seeded in each well of a 6 well plate. The following day, cells were treated with quercetin for approximately 24 hours with DMSO used as a negative control and ActD (Tocris 1229) used as a positive control. Following 24 hours, cells were treated with DRB (5,6-dichlorobenzimidazole 1-β-d-ribofuranoside) (Cayman Chemicals 10010302) at a final concentration of 100 µM and incubated for 3.5 hours. Next, cells were washed with PBS twice and replaced with fresh DMEM supplemented with 10% FBS and 1% penicillin and streptomycin containing 1 µM colchicine or 20 nM ActD or DMSO only, and 5-ethynyl uridine (EU, from the Click-iT Nascent RNA Capture Kit, Thermo Fisher) was added for 30 minutes, 1 hour and 3 hours. Following EU treatment, total cellular RNA was extracted using Quick-RNA Midiprep Kit (Zymo Research Corporation). Nascent RNA was then isolated using the Click-iT Nascent RNA Capture Kit (Thermo Fisher) and cDNA was generated according to the manufacturer’s protocols.

### Fluorescence indicator displacement assay

Target DNA was dissolved in 1X sodium phosphate buffer and annealed by heating at 90°C for 10 min and cooling at room temperature overnight. SYBR Safe (Thermo Fisher) was used as a fluorescent probe and fluorescence (F) was recorded using excitation wavelength at 500 nm and emission wavelength at 530 nm. Each assay solution contained 2 µM DNA, 20 nM SYBR Safe, and a variable amount of quercetin from 0 µM to 100 µM. The percentage of SYBR safe displacement was calculated using equation 4.

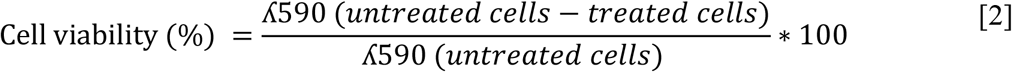

### Computational modeling of quercetin binding to DNA

To gain insights into interaction of quercetin with CTG repeats, a combination of *in silico* modeling and molecular docking was used. First, a 10mer CTG repeat DNA with an intercalation site was modeled based on an ellipticine bound DNA strucutre (PDB ID: 1Z3F) (*40*). Since, ellipticine, is comparable in size to quercetin, we expect their intercalation sites will be comparable as well. Three unique intercaltion sites are possible in a CTG repeat. All three intercalation sites were positioned between the 5^th^ and 6^th^ base pair of the 10mer CTG repeat DNA duplex. The sequence was adjusted accordingly for the three constructs (i) 5′-GCTGC*TGCTG-3′ | 5′-CAGCA*GCAGC-3′ with site CT·AG (ii) 5′-TGCTG*CTGCT-3′ | 5′-AGCAG*CAGCA-3’ with site GC·GC (iii) 5′-CTGCT*GCTGC-3′ | 5′-GCAGC*AGCAG-3’ with site TG·CA; where * denotes the intercalation site. Molecular docking was performed using Molecular Operating Environment (MOE) (https://www.chemcomp.com/Products.htm). First, structure preparation, involved correcting protonation states and topologies of both the DNA and quercetin. Quercetin was modeled with zero net charge at physiological pH (*41*). Next molecular docking was performed using the triangle matcher and london dG method for placement of quercetin and scoring of the complexes respectively, to find the top 50 binding modes within the 3 unique sites. These were then refined using induced fit method to allow for some local flexibility at the intercalation site and scored using GBVI/WSA dG method to obtain the top 5 binding modes. The interaction between quercetin:DNA complex structures were analyzed in MOE and visualized in PyMOL (The PyMOL Molecular Graphics System, Version 2.3.5, Schrödinger, LLC).

### Treatment of fibroblasts

DM1-A, DM2, and unaffected control-A fibroblasts were previously obtained from skin biopsies under a University of Florida-approved Institutional Review Board (IRB) protocol with the informed consent from all subjects and have been described previously (*22*). DM1-B (GM03987) and unaffected control-B (GM08400) fibroblasts were obtained from Coriell Institute. Approximately 1×10^5^ cells were seeded per well of 12 well culture plates and cultured to ∼100 % confluence in DMEM or MEM media containing 15% FBS and 1% antimycotic/antibacterial under standard conditions of 37°C and 5% CO2. Once confluency was reached, cells were washed with PBS and fresh growing media containing quercetin (Sigma-Aldrich Q4951), dasatinib (Sigma-Aldrich SML2589) or A1155463 (Sigma-Aldrich SML3162) dissolved in DMSO was added at the indicated concentrations for 24 hours.

### Treatment of myotubes

Primary patient and unaffected control myoblast cells were previously obtained from biopsies of an individual with DM1 and an unaffected individual (*26*) under a University of Florida-approved Institutional Review Board (IRB) protocol with the informed consent from all subjects. Approximately, 1×10^5^ cells were grown in each well of 12 well culture plates in SkGM-2 BulletKit growth medium (Lonza) to >90% confluency under standard culture conditions of 37°C and 5% CO_2_. Cells were then differentiated to myotubes for 7 days in differentiation medium (DMEM/F-12 50/50 medium (Corning) supplemented with 2% (vol/vol) donor equine serum (HyClone)) under standard culture conditions. After 7 days, cells were washed with PBS and replaced with SkGM-2 BulletKit growth medium (Lonza) containing quercetin dissolved in DMSO at the indicated concentrations for 72 hours under standard culture conditions. Treatments of DM1 myotubes for 24 hours with quercetin was not sufficient to significantly rescue mis-splicing.

### RT-PCR splicing analysis

RT-PCR analysis was carried out as previously described with minor modifications (*15*). Briefly, total cellular RNA was isolated using Quick-RNA Midiprep Kit (Zymo Research Corporation) with on-column DNase I treatment at room temperature for 15-20 minutes. RNA from quadriceps muscle tissue of EMIQ treated and untreated *HSA*^LR^ mice and wildtype mice were TRIZol-extracted. RNA concentration was measured using NanoDrop (Thermo-scientific) and a total of 500 ng of total RNA was used for cDNA synthesis with the SuperScript IV kit (Thermo Fisher) and random hexamer primers (IDT). 2 µL of cDNA was used to perform semi-quantitative PCR using specific primer sets listed in table S2. The resulting PCR products were analyzed by capillary electrophoresis on a Fragment Analyzer with the 1-to 500-bp DNF-905 kit (Advanced Analytical). Pro size 4.0.2.7 software (Advanced Analytical) was used to quantify the electropherogram peaks corresponding to exon inclusion and exclusion products. Percent (%) Exon Inclusion was calculated using equation 1. RFU (Relative Fluorescence Units)

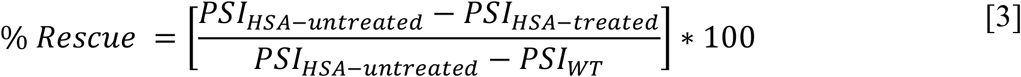

### RT-qPCR analysis in fibroblasts and myotubes

Following drug treatments, total cellular RNA was isolated using Quick-RNA Midiprep Kit (Zymo Research Corporation) with on-column DNase I treatment. RNA concentrations were measured on the Nanodrop (Thermo Fisher) and 500 ng of total RNA from each sample was reverse transcribed into cDNA using SuperScript IV (Thermo Fisher) and random hexamer primers (IDT). qPCR was performed using primers listed in table S3. Data were normalized and analyzed using the ΔΔCt method and plotted in GraphPad Prism (version 9.4.1).

### EMIQ treatment in mice

Mouse handling and experimental procedures were performed in accordance with the Osaka University guidelines for the welfare of animals and were approved by the institutional review board. Homozygous *HSA*^LR^ transgenic mice of line 20b have been described previously (*27*). Age (< 4 months old), gender- and sibling-matched mice were administered enzymatically modified isoquercitrin (EMIQ from San-Ei Gen F.F.I., Inc) in their drinking water at concentrations of 1.5 g/L (low dose) or 15 g/L (high dose) for 6 or 12 weeks. Control mice were drinking tap water throughout the experiments. After the treatments, mice were sacrificed, and the vastus lateralis (quadriceps) muscle was dissected. Total RNA was extracted and cDNA synthesis and PCR amplification were performed as described previously (*42*). Quantitative reverse transcriptase-PCR was performed using TaqMan Gene Expression assays on an ABI PRISM 7900HT Sequence Detection System (Applied Biosystems). The level of endogenous mouse *Dmpk*, and *HSA* transgene-derived mRNA were normalized to *Gtf2b* as described previously (*42*). Electromyography was performed under general anesthesia as described previously(*42*). Briefly, at least 10 needle insertions were performed in the vastus muscle, and myotonic discharges were graded on a four-point scale: 0, no myotonia; 1, occasional myotonic discharge in ≤ 50% of needle insertions; 2, myotonic discharge in > 50% of insertions; and 3, myotonic discharge with nearly all insertions.

### Immunofluorescence

Frozen vastus lateralis muscle from *HSA*^LR^ mice was sectioned into 10 μm slices onto slides and fixed with 10% (v/v) buffered formalin, permeabilized using 1:1 methanol/acetone and blocked using Background Sniper (Biocare Medical). Slides were incubated with 1:100 rabbit anti Clcn1 (Alpha Diagnostic International) overnight at 4 °C. Samples were incubated with 1:1000 goat antirabbit Alexa Fluor 488 (Thermo Fisher) for 1 h at RT and mounted using VECTASHIELD Hard·Set Mounting Medium with DAPI (Vector Laboratories). Samples were imaged on BZ-X810 fluorescent microscope (Keyence).

### RNAseq library preparation

The library preparation was performed as previously described with minor modifications (*15*). Briefly, total RNA was TRIZol-extracted from the quadricep muscle of wild-type FVB mice and EMIQ-treated/untreated mice. RNA quality was assured by using capillary electrophoresis on a fragment analyzer (Advanced Analytical). DNF 474 kit (Advanced Analytical) was used to run the RNA sample and RIN number >8 was used to assure the quality of RNA. The NEBNext UltraII Directional RNA Library Prep Kit for Illumina (NEB #E7760) with NEBNext rRNA Depletion Kit v2 (NEB #E7400) was used to prepare RNAseq libraries, with a total of 1000ng of RNA input for each library. The manufacturer’s protocol was followed, with the following exceptions: 40X adapter dilutions were used and 8 cycles of library amplification were performed. The resulting libraries underwent quality control. The average fragment size was calculated via capillary electrophoresis on the Fragment Analyzer (Agilent) using the DNF-474 High Sensitivity Next Generation Sequencing Fragment assay. Library concentration was quantified using the Qubit dsDNA High Sensitivity Assay (ThermoFisher Scientific #Q32851) and with the NEBNext Library Quant Kit for Illumina (NEB #E7630). Libraries were pooled in equimolar amounts to a final concentration of 2 nM and were sequenced using paired-end, 100 base pair sequencing on the Illumina NextSeq 2000 at The RNA Institute, SUNY at Albany.

### Gene expression analysis of RNAseq data

The gene expression analysis was performed as previously described (*15*) with minor modifications. Briefly, raw reads were aligned to the GRCm39.105 mouse genome using STAR (Version 2.7.5). Uniquely aligned paired sequences were used as input for Stringtie (version 1.3.4d) and prepDE.py was used to generate gene counts. DESeq2 (version 1.3.4d) was used to perform differential gene expression analysis. Differential gene expression was considered significant where adjusted *P* < 0.1.

### Splicing analysis of RNAseq data

Uniquely aligning paired sequences were input to rMATS turbo (v4.1.1) to analyze the isoform abundance and compared to wild-type mouse sequence (*15*). Cassette exon events were considered significant where FDR < 0.05 and *P* < 0.0005. Percent splicing rescue was calculated using equation 3. Where PSI is the percent exon inclusion explained in equation 1. *HSA*-treated and *HSA*-untreated are *HSA*^LR^ mice treated with EMIQ and untreated respectively.

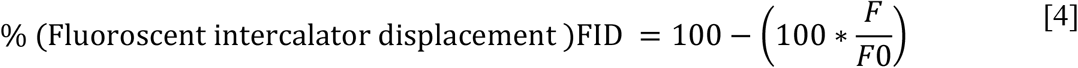

Of those events, a percent rescue of more than 10% and less than 110% was considered a “Rescue” with EMIQ treatment.

### Quantification and statistical analysis

Data are expressed as mean ± standard deviation. All data shown are the summary of three or more biological replicates and statistical analyses were completed in Prism 7. For data sets where treatments were compared individually to a control, two-tailed student’s t-test was used to determine statistical significance with the associated *P* value. When multiple groups were compared to a control, one-way ANOVA was used with Dunnett’s multiple comparisons test. Statistical significance values used: \**P* < 0.05, \*\**P* < 0.01, \*\*\**P* < 0.001, \*\*\*\**P* < 0.0001.

## Acknowledgments

Special thanks to members of the Berglund, Reddy and RNA Institute labs for helpful discussions, experimental advice, and comments on the manuscript.

## Funding

These studies were supported by funding from the NIH (P50NS048843, R01NS120485, R01NS104010) and startup funds from the University at Albany to J.A.B. and K.R.; Intramural Research Grant (2-5) for Neurological and Psychiatric Disorders of National Center of Neurology and Psychiatry to M.N.; Myotonic Dystrophy Foundation postdoctoral fellowship to S.K.M.; Myotonic Dystrophy Foundation predoctoral fellowship to J.A.F. and a NIH T32 training fellowship (5T32GM132066) to S.M.H.

## Author contributions

S.K.M, S.M.H, J.A.F, S.V. and M.N. conducted experiments; all authors contributed to data analysis and interpretation; S.K.M, K.R. and J.A.B. wrote the manuscript with input from all authors; J.D.C, K.R. and J.A.B. supervised the research.

## Competing interests

S.K.M., J.D.C., K.R. and J.A.B. have filed a provisional patent application for the use of quercetin and related flavonoids for the treatment of myotonic dystrophy. J.A.B. serves as a consultant for Entrada Therapeutics, Kate Therapeutics, Juvena Therapeutics and Syros Pharmaceuticals.

## Data and materials availability

RNA-seq data have been deposited in the Sequence Read Archive (SRA) database (BioProject accession number PRJNA891268), https://www.ncbi.nlm.nih.gov/sra.

## Notes

https://www.ncbi.nlm.nih.gov/sra.

