## Supplementary Material for "Quercetin selectively reduces expanded repeat RNA levels in models of myotonic dystrophy"

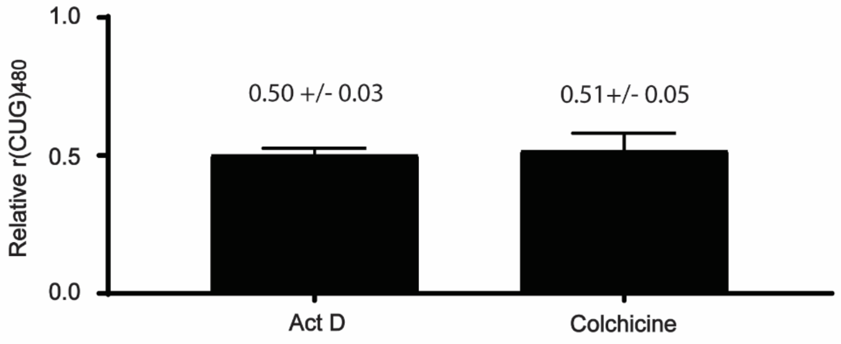


**fig. S1 Positive controls from CTG repeat-selective HeLa DM1 screen of NCI Natural Compound Set V.** Treatment of HeLa DM1 cells with Actinomycin D (ActD) at 20 nM and colchicine at 1µM for 24 hours in triplicate across the library. Multiplex RT-qPCR was performed to measure $r(CUG)_{480}$ levels relative to $r(CUG)_0$. (Mean $\pm$ SD, n = 21 treatments).

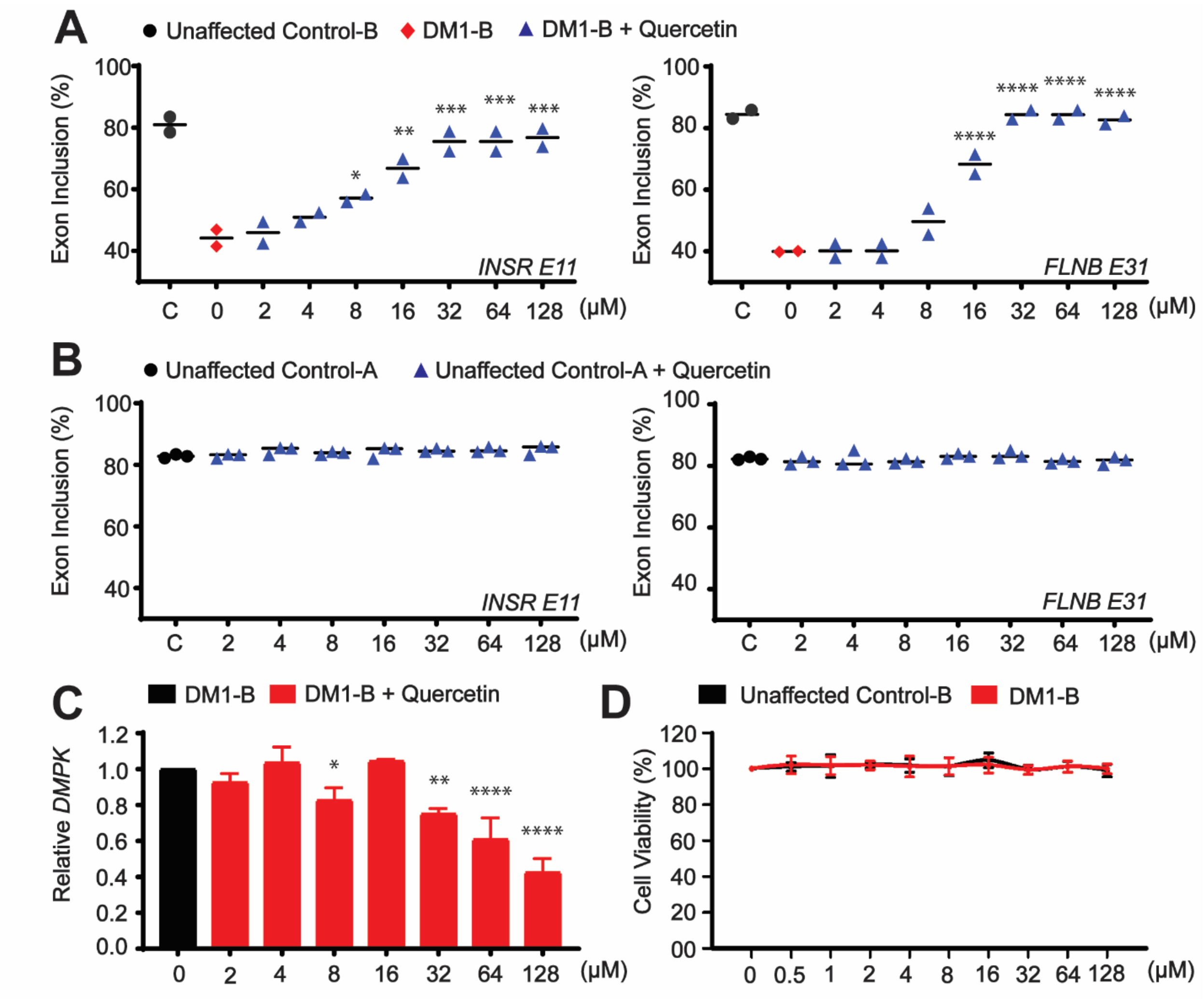


**fig. S2 Validation of quercetin activity in additional DM1 patient fibroblasts. (A)** RT-PCR cassette exon isoform analysis of *INSR exon 11* (left) and *FLNB exon 31* (right) alternative splicing events following 24-hour treatment with quercetin (Mean ± SD, n = 2 biological replicates). **(B)** RT-PCR cassette exon isoform analysis of *INSR exon 11* (left) and *FLNB exon 31* (right) alternative splicing events following 24-hour treatment with quercetin (Mean ± SD, n = 3 biological replicates). (**C**) RT-qPCR analysis of *DMPK* levels (relative to *GAPDH*) following quercetin treatment normalized to DMSO control (0μM) in DM1 fibroblasts (Mean ± SD, n = 2 biological replicates. (**D**) Cell viability assay performed on unaffected control and DM1 fibroblasts following 24-hour treatment with quercetin at the indicated concentrations. Normalized to DMSO control (0 μM) (Mean ± SD, n = 2 biological replicates). One-way ANOVA with Dunnett's multiple comparisons test comparing treated fibroblasts to DMSO control (0μM). *$P < 0.05$, **$P < 0.01$, ***$P < 0.001$ ****$P < 0.0001$.

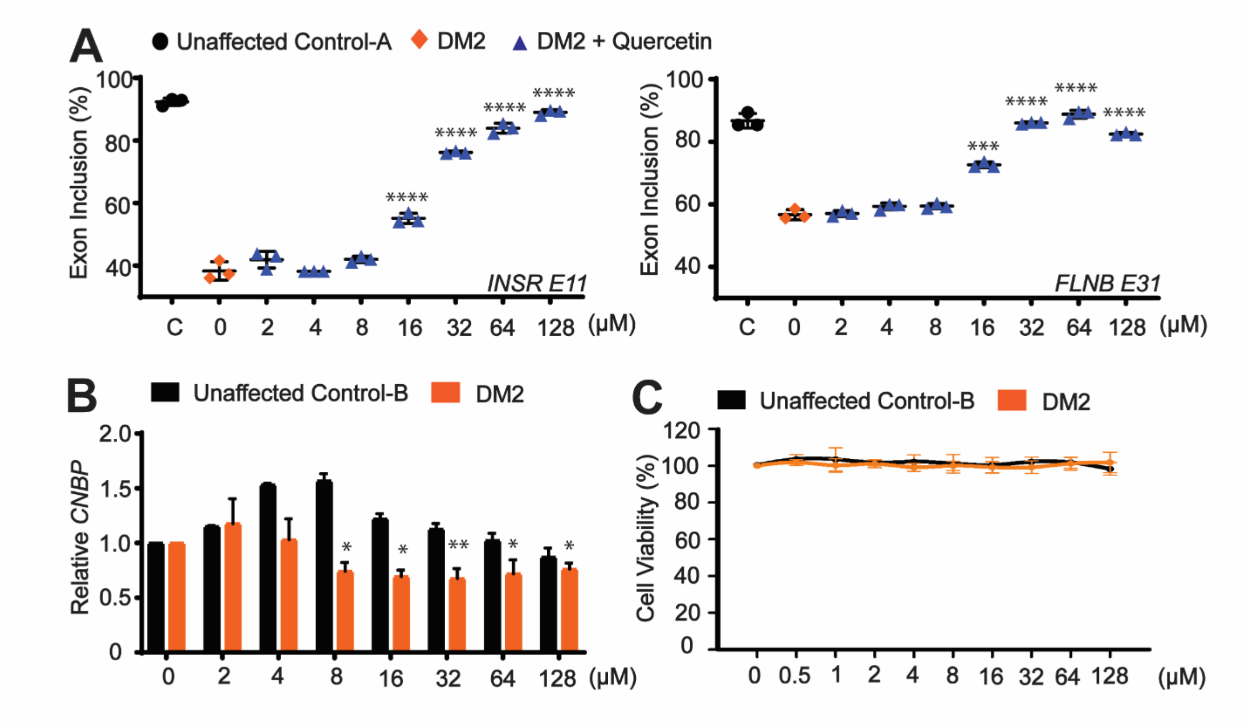


**fig. S3 Validation of quercetin activity in DM2 patient derived fibroblasts.** (**A**) RT-PCR cassette exon isoform analysis of *INSR exon 11* (left) and *FLNB exon 31* (right) alternative splicing events following 24-hour treatment with quercetin (Mean ± SD *n* = 3 biological replicates). (**B**) RT-qPCR analysis of *CNBP* levels (relative to *GAPDH*) following quercetin treatment normalized to DMSO control in DM2 and unaffected control fibroblasts (Mean ± SD *n* = 3 biological replicates). (**C**) Cell viability assay performed on unaffected control and DM2 fibroblasts following 24-hour treatment with quercetin at the indicated concentrations. Normalized to DMSO control (0 µM) (Mean ± SD, n = 3 biological replicates). One-way ANOVA with Dunnett's multiple comparisons test comparing treated fibroblasts to DMSO control (0µM). *$P < 0.05$, **$P < 0.01$, ***$P < 0.001$, ****$P < 0.0001$.

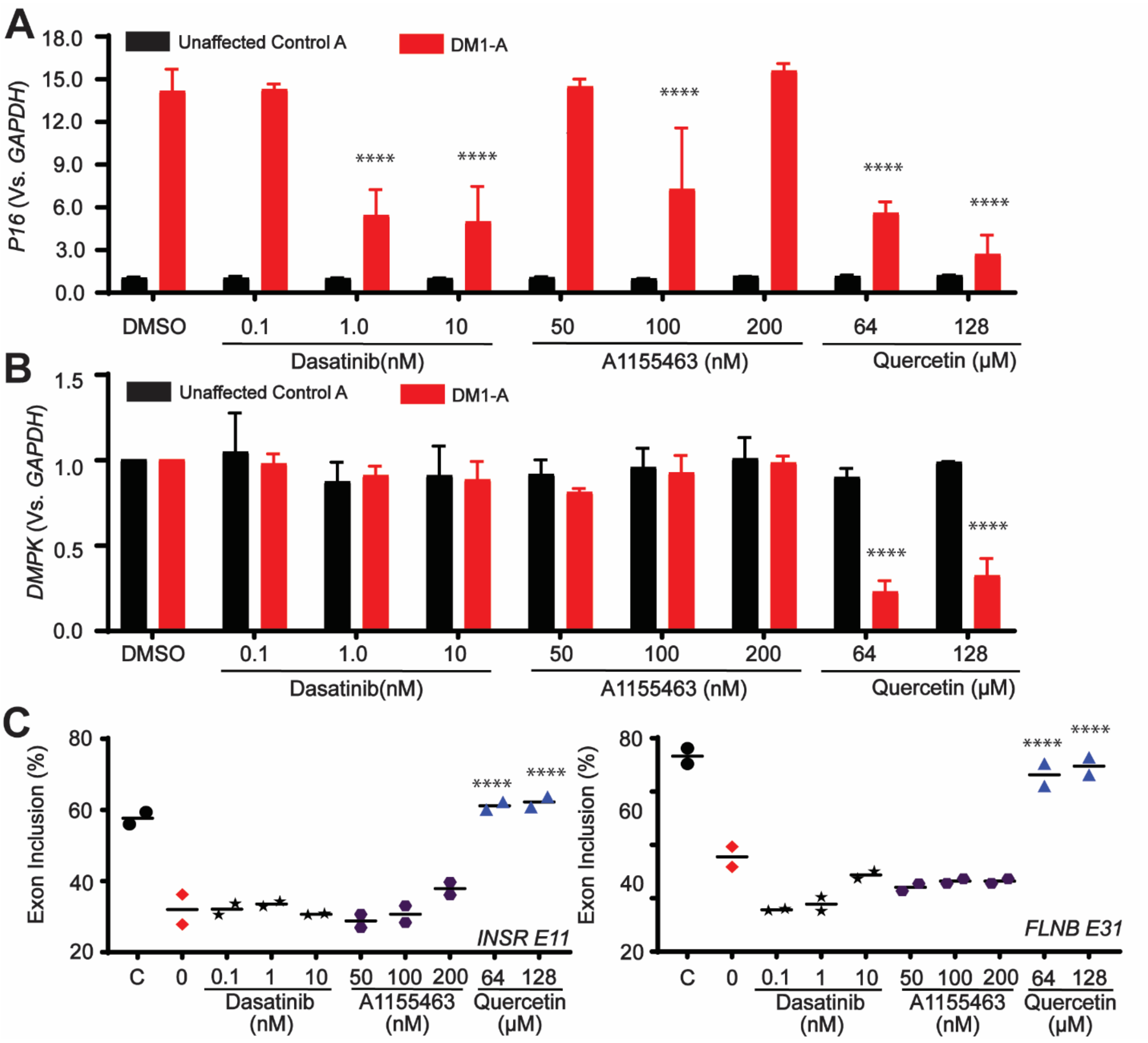


**fig. S4 Effects of senolytics in DM1 patient fibroblasts.** (**A**) RT-qPCR analysis of *p16* senescence marker expression levels (relative to *GAPDH*) following treatment unaffected control and DM1 patient fibroblasts with dasatinib, A1155463 and quercetin normalized to DMSO control (0µM) for 24 hours (Mean ± SD, n = 2 biological replicates). (**B**) RT-qPCR analysis of *DMPK* levels (relative to *GAPDH*) following dasatinib, A1155463 or quercetin treatments normalized to DMSO control (0µM) in DM2 fibroblasts (Mean ± SD, n = 2 biological replicates). (**C**) RT-PCR cassette exon isoform analysis of *INSR exon 11* (left) and *FLNB exon 31* (right) alternative splicing events following 24-hour treatment with dasatinib, A1155463 or quercetin in DM1 fibroblasts. (Mean ± SD, n = 2 biological replicates). One-way ANOVA with Dunnett's multiple comparisons test comparing treated fibroblasts to DMSO control (0µM). ****$P < 0.0001$.

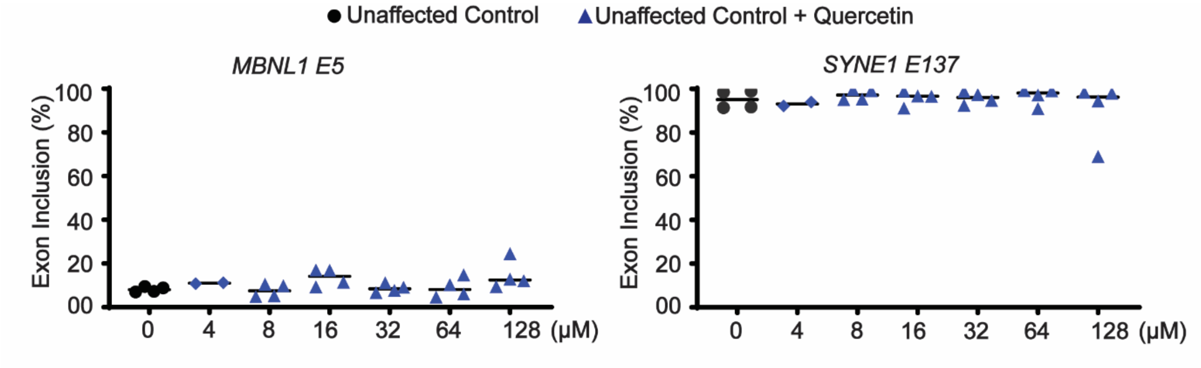


**fig. S5 Treatment of unaffected control myotubes with quercetin.** RT-PCR cassette exon isoform analysis of *MBNL1 exon 5* (left) and *SYNE1 exon 137* (right) alternative splicing events following 72-hour treatment with quercetin at the indicated concentrations (Mean ± SD, n = 4 biological replicates).

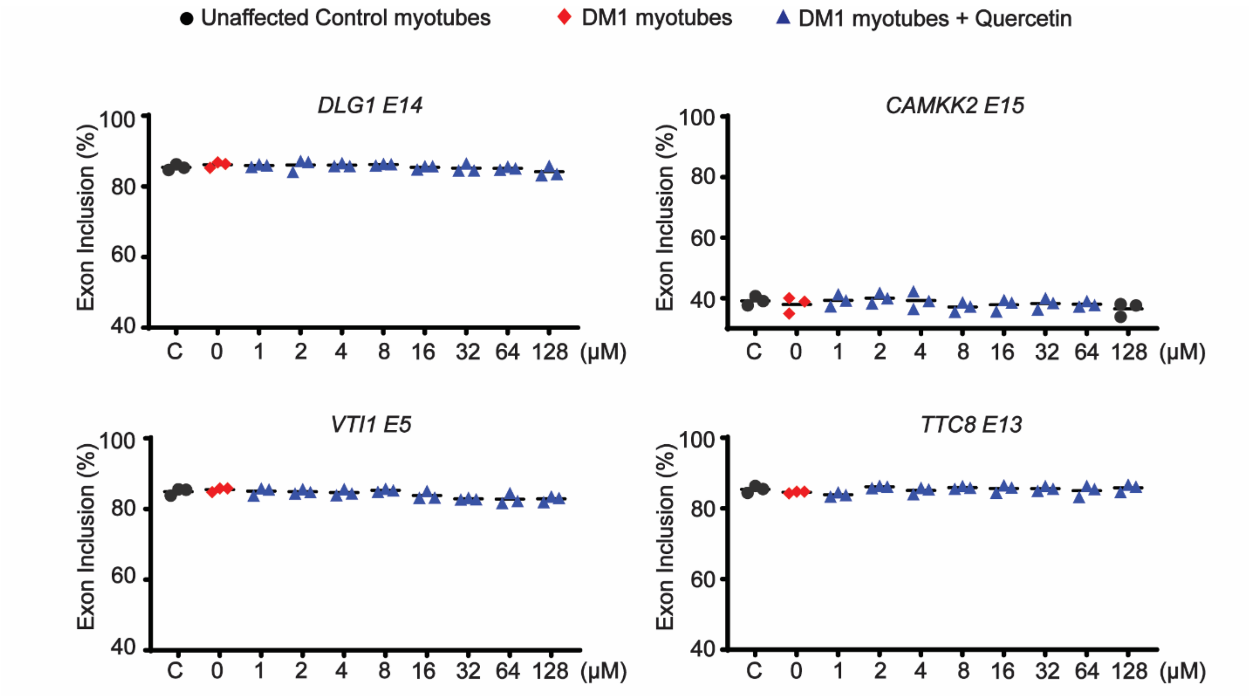


**fig. S6 Evaluation of non-DM1 associated alternative splicing events in DM1 myotubes treated with quercetin.** RT-PCR cassette exon isoform analysis of the indicated alternative splicing events following 72-hour treatment with quercetin at the indicated concentrations (Mean ± SD, n = 3 biological replicates).

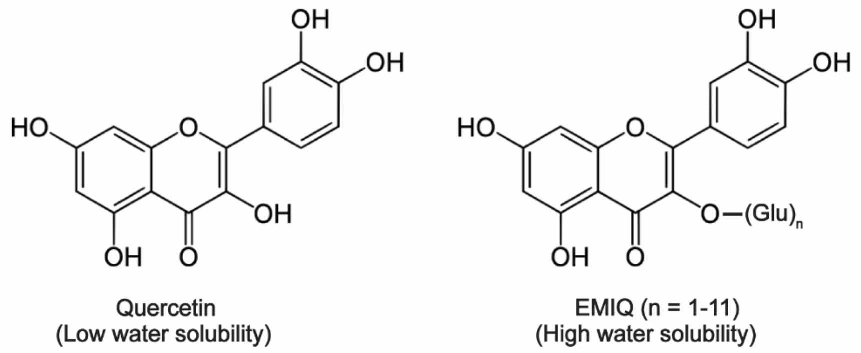


**fig. S7 Structures of quercetin and enzymatically modified isoquercitrin (EMIQ).** While quercetin has poor water solubility and bioavailability, the addition of glucose moieties to form EMIQ increases solubility and boosts bioavailability. EMIQ is processed by α-glucosidase and subsequently by lactase-phlorizin hydrolase to quercetin which is absorbed through the small intestinal epithelium. (www.saneigen.com).

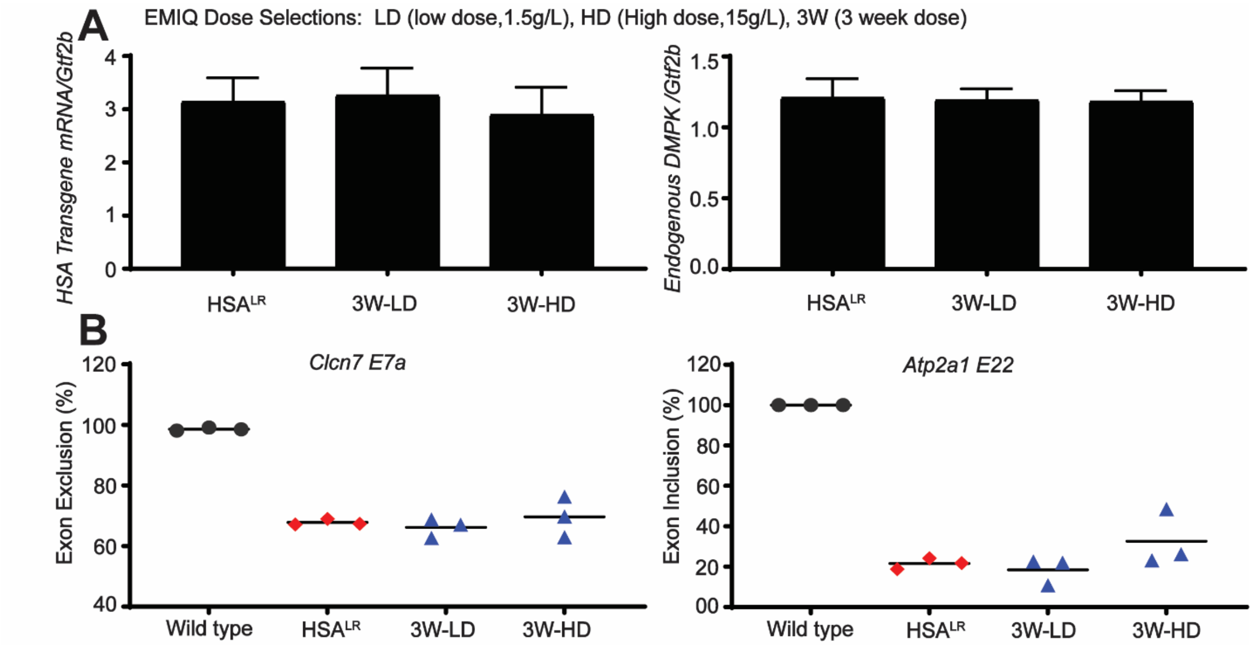


**fig. S8 Treatment of *HSA*$^{LR}$ DM1 transgenic mouse model with EMIQ for 3 weeks.** (**A**) RT-qPCR analysis of *HSA* transgene (left panel) or endogenous *Dmpk* (right panel) expression (normalized to *Gtf2b*) following EMIQ administration in the drinking water for 3 weeks (3W) at a low dose of 1.5 g/L (LD) or high dose of 15 g/L (HD). Mean ± SD, n = 3 biological replicates. (**B**) RT-PCR cassette exon isoform analysis of *Clcn1 exon 7a* (left) and *Atp2a1 exon 22* (right) alternative splicing events following EMIQ administration in the drinking water at the indicated doses. Mean ± SD, n = 3 biological replicates.

### Primers and probes

##### Table S1. Primers/probes used for HeLa DM1 cellular screen

| Name | Sequence |
|---|---|
| HT_RT | 5´-CTACACGACGCTCTTCCGATCTTCTTATCGAATGTCGGGGTCTCAGTGC-3´ |
| HT_Forward | 5´-CGATCTCTGCCTGCTTACTC |
| HT_Reverse | 5´-GTCGGAGGACGAGGTCAATAAA |
| HT_Probe1 | /56FAM/AGAGCAGCG/ZEN/CAAGTGAGGAGG/3IABkFQ/ |
| HT_Probe2 | /5HEX/TGACGCAGC/ZEN/CACGTGAAGGTC/3IABkFQ/ |

##### Table S2. Primers used for RT-PCR splicing assay

| Target | Forward Primer | Reverse Primer |
|---|---|---|
| *Human INSR exon 11* | 5´- CCTGTCCAAAGACAGACTCTCAGATCCTG | 5´- GTCGAGGAAGTGTTGGGGAAAGC |
| *Human FLNB exon 31* | 5´- GCTTCGGTGGTGTTGATATTC | 5´- GTCACTCACTGGGACATAGG |
| *Human MBNL1 exon5* | 5´- AGGGAGATGCTCTCGGGAAAAGTG | 5´- GTTGGCTAGAGCCTGTTGGTATTGG |
| *Human MBNL2 exon5* | 5´- ACAAGTGACAACACCGTAACCG | 5´- TTTGGTAAAGGATGAAGAGCACC |
| *Human SYNE1 exon137* | 5´- GACAAAGATTTCTACCTCCGGGG | 5´- CCCAGTTGTCGGATCTGTGACTC |
| *Mouse Atp2a1 exon22* | 5´- GCTCATGGTCCTCAAGATCTCAC | 5´- GGGTCAGTGCCTCAGCTTTG |
| *Mouse Clcn1 exon7a* | 5´- TGAAGGAATACCTCACACTCAAGG | 5´- CACGGAACACAAAGGCACTG |
| *Human CAMKK2 exon15* | 5’- CCTGGTGAAGACCATGATACG | 5’- GGCCCAGCAACTTTCCAC |
| *Human DLG1 exon14* | 5’- AGCCCGATTAAAAACAGTGAAA | 5’- CGTATTCTTCTTGACCACGGTA |
| *Human TTC8 exon3* | 5’- AGCTATTTTAGGCGCAGGAAGT | 5’- CATTTTCATCCAGCATCATTTCTG |

##### Table S3. Primer used for qPCR

| Target | Forward Primer | Reverse Primer |
|---|---|---|
| Human *DMPK* | 5´- CACGTTTTGGATGCACTGAGAC | 5´- GATGGAGGGCCTTTTATTCGCG |
| Human *GAPDH* | 5´- AATCCCATCACCATCTTCCA | 5´- TGGACTCCACGACGTACTCA |
| Human *CNBP* | 5´- TACCTTGCGAGCCGTCTTC | 5´- CACTCATTGCTGCTCATGG |
| Human *p16* | 5´- TGTTCGCATTGCCAAGGTTC | 5´- CGTTTCCGTGAATGTTGTCCC |
| Human *p21* | 5´- CTGGGGATGTCCGTCAGAAC | 5´- GTGACAGGTCCACATGGTCT |
